# Reliable multiplex generation of pooled induced pluripotent stem cells for genetic testing

**DOI:** 10.1101/2022.08.10.500520

**Authors:** Molly Smullen, Julia M Reichert, Pepper Dawes, Qi Wang, Benjamin Readhead, George M Church, Elaine T Lim, Yingleong Chan

## Abstract

Inducing somatic cells into pluripotent stem cells (iPSCs) provides an excellent model for studying systems *in-vitro*. Understanding the impact of individual donor genetic backgrounds on reprogramming ability would allow researchers to harness these genetic differences and increase the efficiency of the reprogramming process. To better understand the genetic basis of reprogramming cells into iPSCs, we present Induction of Pluripotency from Pooled Cells (iPPC) - an efficient, scalable, and reliable reprogramming procedure. Using our deconvolution algorithm that employs low-coverage pooled sequencing and single nucleotide polymorphisms (SNPs), we estimate individual donor proportions of cell lines within large cohorts. With iPPC, we concurrently reprogrammed over one hundred donor LCLs into iPSCs and found strong correlations of individual donors’ reprogramming ability across multiple experiments. We note that individual donors’ reprogramming ability remains consistent across both same-day replicates and multiple experimental runs, and that the expression of certain immunoglobulin precursor genes (IGLV10-54, IGLV3-9, IGLV1-17, IGLV1-6, and IGLV3-1) may impact reprogramming ability. Our process enables a multiplex framework to study the reprogramming ability of different donor cells into iPSCs and also provides a reliable method along with a pooled library of donor iPSCs for downstream research and investigation of other *in-vitro* phenotypes.

## Introduction

Several years ago, Takahashi et. al reported the ability to transform somatic fibroblasts into induced pluripotent stem cells (iPSCs)(Takahashi et al., 2007). Such iPSCs can be generated by expressing a few transcription factors in the adult tissue, principally a subset of OCT4, SOX2, KLF4, MYC, LIN28 and NANOG(Takahashi et al., 2007; Yu et al., 2007). This discovery has since enabled researchers to turn almost any adult somatic tissue type from any donor individual into iPSCs for the purpose of studying complex diseases, drug development and to advance the development of personalized medicine(Chang et al., 2020; Rowe and Daley, 2019; Silva and Haggarty, 2020). These iPSCs offer an alternative to model organisms and other *in-vitro* systems as they can be differentiated into a wide variety of tissue types for many scientific applications such as *in-vitro* patient-specific disease modeling and drug testing(Paik et al., 2020; Qian and Tcw, 2021; Lim et al., 2022).

Somatic cells must overcome a number of barriers to be successfully reprogrammed into iPSCs(Haridhasapavalan et al., 2020). It was first reported that direct transcription factor reprogramming had success rates between 0.02% to 2%(D et al., 2008; Maherali et al., 2008; Takahashi et al., 2007; Yu et al., 2007). As an attempt to optimize reprogramming efficiency, researchers have tried developing alternative reprogramming methods, using tools like CRISPR/Cas9, episomal vectors, miRNA, and small molecules(Zeng et al., 2018; Liu et al., 2020; Chen et al., 2020). Others have uncovered information about the changes that cells undergo and what factors play a key role in the transition from a somatic to stem cell state in the attempt to better understand what affects the reprogramming process(Mahmoudi et al., 2019; Zhou et al., 2019; Karagiannis et al., 2019; Ray et al., 2021).

The variability between different donor genetic backgrounds could also affect the efficiency to which cells can be reprogrammed into iPSCs and the reasons underlying the variability could be exploited for increasing reprogramming efficiency. In one study, the authors showed that the contribution of genetics in post-reprogramming iPSC traits can be as high as 45%, which suggests that perhaps genetics can also strongly contribute to reprogramming ability(Kilpinen et al., 2017). However, conducting a large-scale study to test the reprogramming ability of a population of donor cells would be a time and resource intensive process. This is compounded further by the need for a genome-wide genetic profile for each donor, which is costly to generate(Schwarze et al., 2018).

Here, we describe our approach for performing such a study which overcomes some of the challenges mentioned. Our approach generates iPSCs from many different donor lymphoblastoid cell lines (LCLs) from the Harvard Personal Genome Project (PGP). The PGP consists of participants with high coverage whole-genome sequencing (WGS) data, self-reported phenotypic data and LCLs available to scientists for research(Ball et al., 2012, 2014; Chan et al., 2017; Mao et al., 2016). With LCLs, it has been demonstrated that they can be successfully reprogrammed into iPSCs via nucleofection of reprogramming factors(Barrett et al., 2014; Choi et al., 2011; Kumar et al., 2016; Muñoz-López et al., 2016; Rajesh et al., 2011; Thomas et al., 2015). In Kumar et. al., the authors reported that LCLs from over 200 different donors can be individually reprogrammed into iPSCs successfully(Kumar et al., 2020). As such, in this manuscript, we describe our method, Induction of Pluripotency from Pooled Cells (iPPC), which is our multiplexed approach to perform pooled reprogramming of multi-donor cells into iPSCs.

The use of iPPC has several advantages over individual donor cell reprogramming. First, it enables the multiplex generation of iPSCs from many different donors within a single experiment thereby saving both resource and time when compared to generating the iPSCs for each donor individually. Next, it allows for a consistent extracellular environment for all the different donor cells which will minimize phenotypic differences due to environmental variability when interrogating the phenotypic effect of genetic variants. Finally, for experiments that make comparisons between different donors within the mixed pool, iPPC is robust to errors caused by sample mix-up and experimental variation across different samples. While iPPC has these advantages, it also introduces its own challenges. First, there is a challenge of selecting only high-quality reprogrammed clones that are pluripotent. While it has been shown that this can be overcome by performing fluorescent or magnetic activated cell sorting to sort for cell expressing pluripotent markers, we demonstrated that we can achieve a high purity of pluripotent cells using magnetic activated cell sorting(Kahler et al., 2013; Yang et al., 2015). Then, there is the challenge of discerning the quantity of each individual donor cell population within the mixed pool and this is not a trivial problem as the cells are morphologically identical to one another even though they are from different donors.

In order to separate out and obtain individual donor proportions from the mixed pool, we utilized our previously published method to quantify such proportions from pooled short read DNA sequencing(Chan et al., 2018). The method does not require exogenous barcodes or single-cell sequencing but employs an algorithm that incorporates single-nucleotide polymorphisms (SNPs) inherent for each donor and uses them as naturally occurring barcodes to track individual donor proportions within the pool of cells. While whole-genome genotypes have to be available for every individual donor as a prerequisite, the method requires only low-pass whole-genome short read sequencing such that a single-read coverage over half a million SNP sites would be sufficient to accurately estimate the proportions of a mixed pool of 100 donors(Chan et al., 2018). By performing iPPC, we accurately measured the individual donor proportions from the iPSC pools generated from pooled reprogramming of the 120 donor LCLs. By comparing the donor proportions for each pool, we determined an inherent reprogramming success phenotype for each donor individual. We found that almost all of our donor LCLs were successfully reprogrammed into iPSCs and the reprogramming success phenotype for each donor is extremely consistent between different experimental replicates, with Pearson’s correlation coefficient estimates over 90%. Due to the high consistency between replicates, we questioned whether there may be any genetic or population factors that would be associated with reprogramming success. We therefore performed a genome-wide association study but did not observe any SNPs with genome-wide significance. We also tested for association with sex and age and did not observe any significant associations. Finally, we looked at the pre-reprogramming cell state by testing for LCL gene expression with the reprogramming success phenotype. To that end, we did observe that the expression of several immunoglobulin lambda variable genes in LCLs were significantly associated with the reprogramming success phenotype. Overall, we demonstrated that iPPC can be used to consistently generate pooled libraries of different donor iPSCs as well as evaluate different reprogramming rates between different donor LCLs.

## Results

### Successful reprogramming of most donor LCLs into iPSCs

A cohort of 120 LCLs from the Harvard Personal Genome Project (PGP) were pooled in equal proportions before nucleofecting three same-day replicates with plasmids expressing hSOX2, hKLF4, OCT3/4, LIN28, c-Myc, and shp53 (**Figure 1**) (See Materials and Methods). We found that by 14 days post nucleofection, a subset of cells had transitioned from culturing in suspension to physically adhering to a Matrigel coated plate(Kumar et al., 2020). We subsequently performed magnetic bead purification to enrich for pluripotent stem cells (See Materials and Methods) and evaluated the pluripotency of the adherent cells using stem cell markers TRA-1-60 and SSEA-4 with flow cytometry, which showed average positive staining of 98.2% and 100% respectively across 3 experimental runs (**Figure 2**). These results show that we have successfully generated iPSCs from a pooled culture of donor LCLs and that we can enrich for purity using magnetic bead selection.

**Figure 1:**
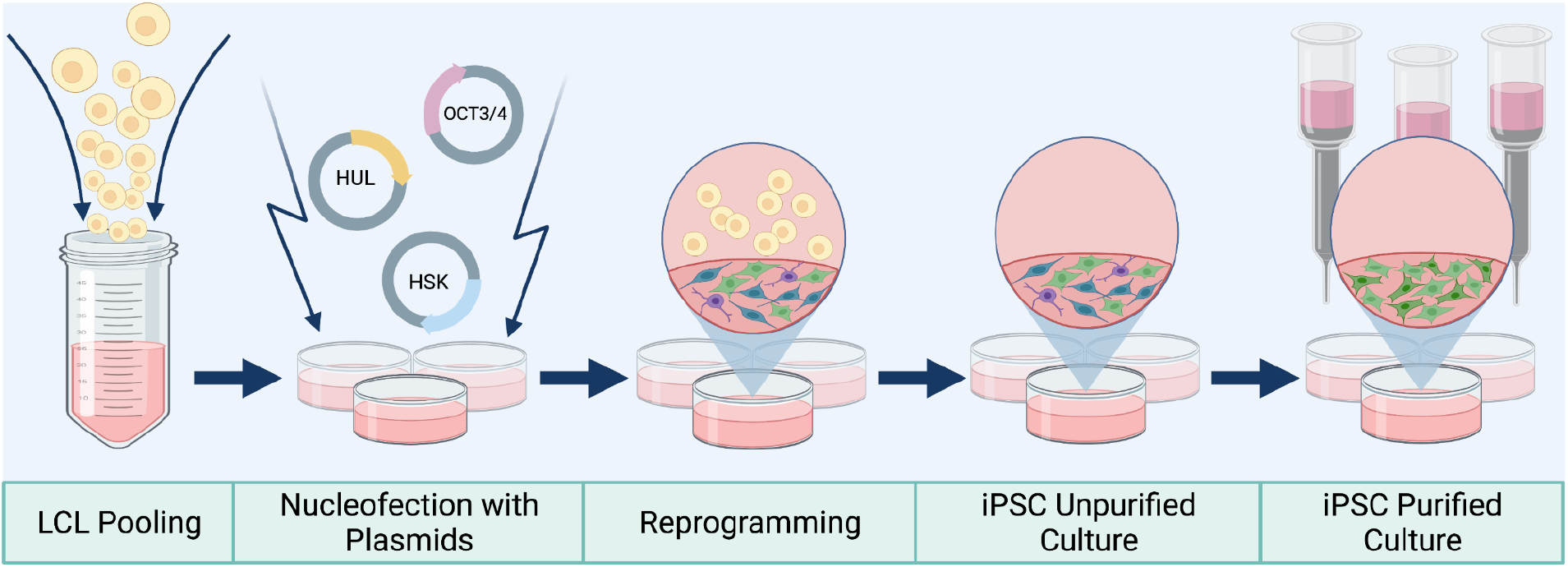
Induction of Pluripotency from Pooled Cells (iPPC) pipeline The figure depicts our experimental method for generating iPSCs from a pooled sample of LCLs. Pooled cells were nucleofected with three plasmids, followed by culture in reprogramming media. Suspended cells were washed away and resulting adherent cells were purified using magnetic bead columns.

**Figure 2:**
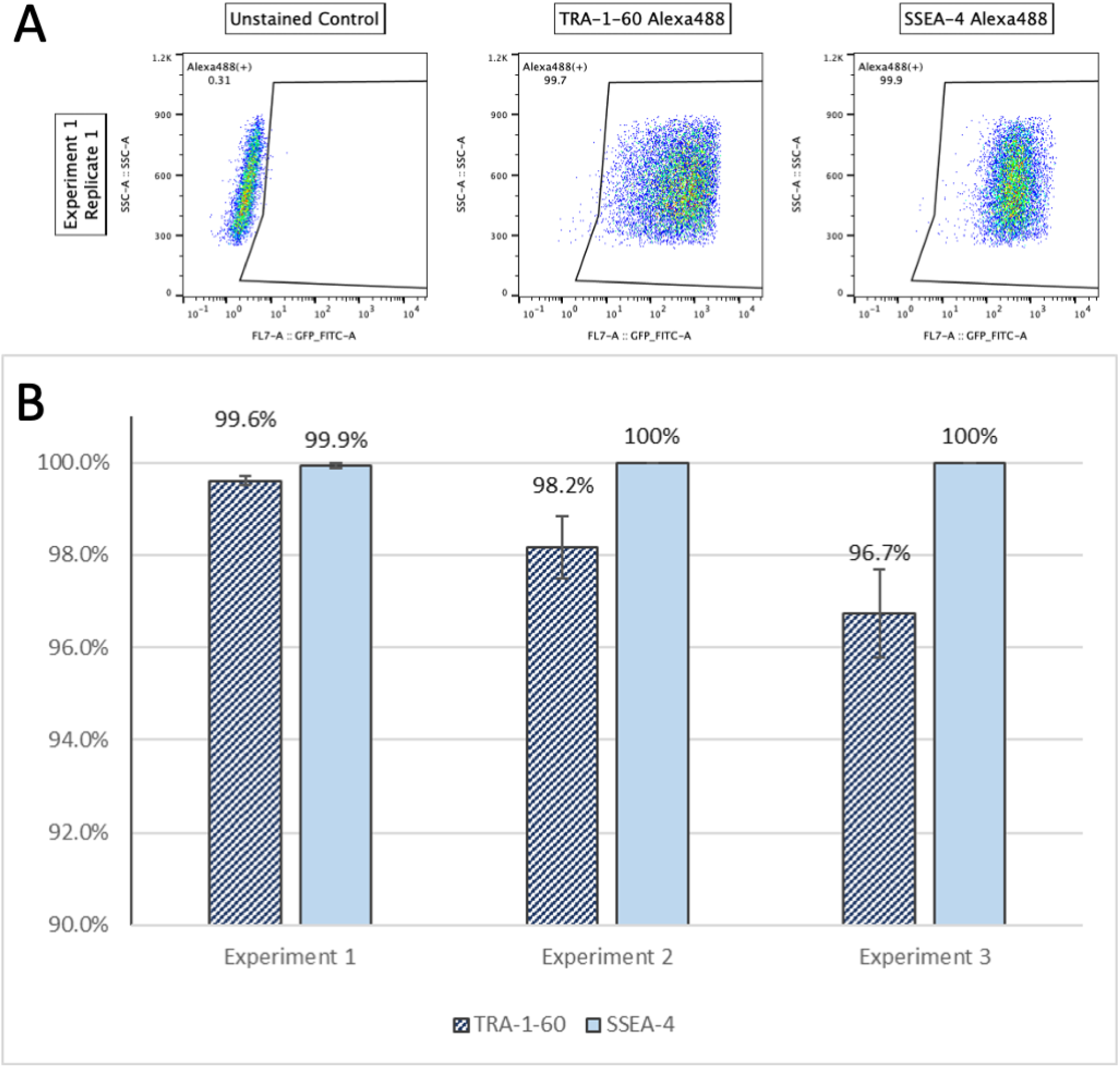
Staining of cells for markers of pluripotency (**A**) Flow cytometry scatter plots from Exp.1-1 purified iPSCs. All plots have the green FITC channel on the x-axis and side scatter on the y-axis. The plot on the left shows an unstained control sample which was used to gate for positive Alexa488 staining. Percent positive staining shown in the top left corner of each plot. The middle and right plots show the TRA-1-60 Alexa488 and SSEA-4 Alexa488 staining respectively. (**B**) Bar chart depicting the average staining of pluripotency markers across from the 3 replicates of each experiment with standard deviation bars. Standard deviations of TRA-1-60 staining for Exp.1, Exp.2, and Exp.3 were 0.1, 0.7, and 1.0% respectively. Standard deviations of SSEA-4 staining for Exp.1, Exp.2, and Exp.3 were 0.1, 0.0, and 0.0% respectively.

### Consistent reprogramming variability observed between different donors

To evaluate the reprogramming ability for each donor, we performed low coverage whole genome sequencing using DNA extracted from the purified iPSC pool and estimated individual donor proportions(Chan et al., 2018). For the initial analyses, we estimated donor proportions by including an additional 20 donors from the PGP cohort where cells were not present in any of the pools to serve as negative controls for our deconvolution method (**Table S1**). The method uses an expectation maximization algorithm to estimate individual donor proportions within the pool from the sequenced DNA (See Materials and Methods). We find that the donor proportions estimates converge to stable estimates after about 100 iterations (**Figure 3**).

**Figure 3:**
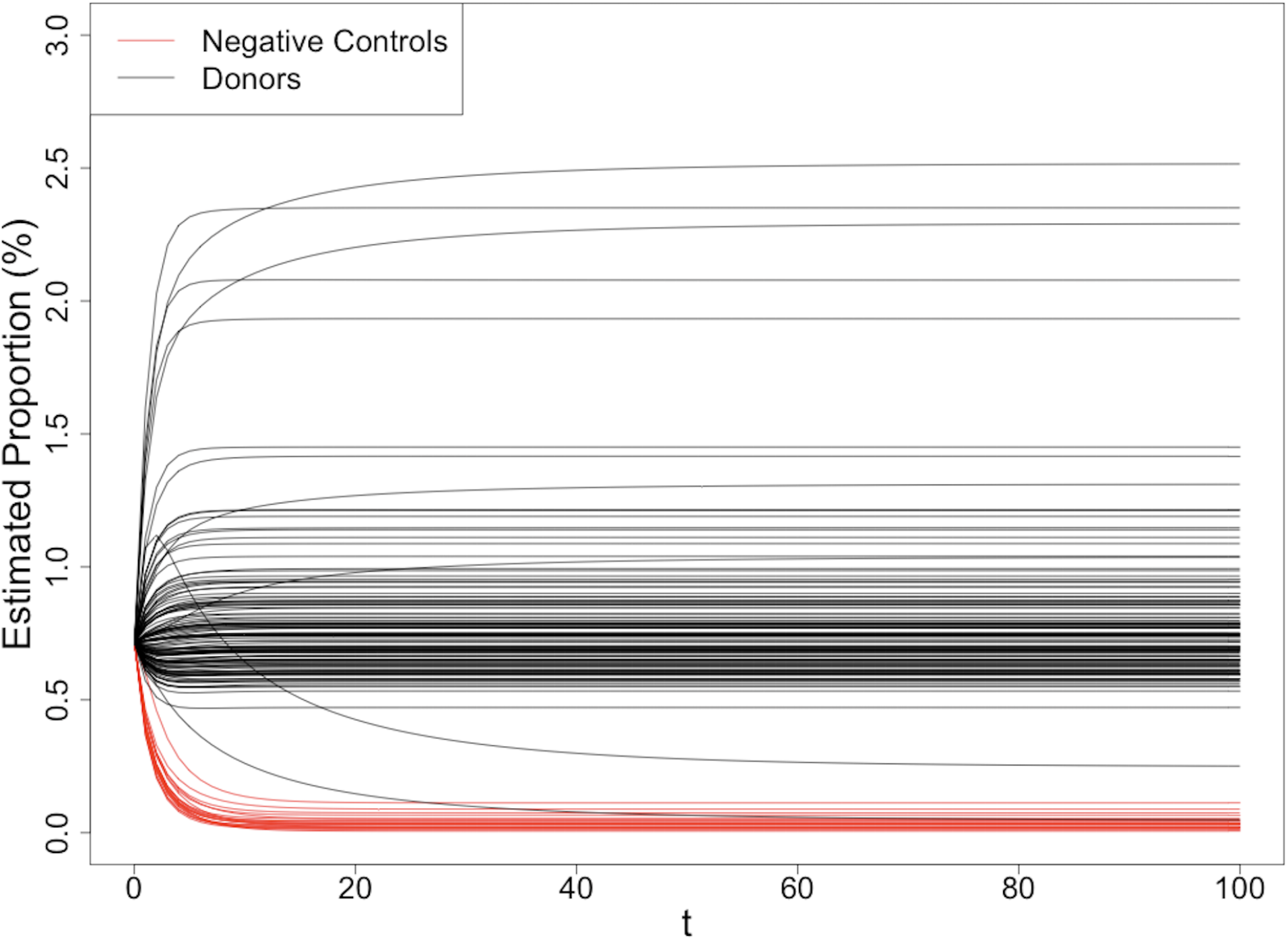
Donor proportion estimate trajectories for Exp.1-1 The figure shows the trajectories of the estimated proportion after each iteration of the algorithm for the data generated for Exp.1-1. The horizontal axis represents the iteration count (t) and the vertical axis represents the estimated proportion. The red trajectories indicate the 20 donors where their cells were absent and they serve as negative controls.

Our deconvolution algorithm returned extremely low proportion estimates (≤0.25%) for each of these 20 donors (**Table S1**). Using the maximum estimated proportion returned for a negative control, we selected 0.28% as the cutoff for successful reprogramming. Individuals with an estimated proportion greater than 0.28% were considered successfully reprogrammed, and those with an estimated proportion less than or equal to 0.28% were considered not successfully reprogrammed (**Table S1**). Based on these results, 118 out of the 120 individuals actually present in our sample were successfully reprogrammed (see Materials and Methods).We recomputed the proportion estimates using only these 118 donors and used the new estimates for downstream analysis (**Table S2**).

Using the new proportion estimates from the final pool of 118 successfully reprogrammed donors, we analyzed the spread of the three same-day replicates from each experiment. Assuming an even distribution of individuals the final reprogrammed iPSC pool, we would expect each individual’s estimated proportion in the final pool to be 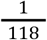, or 0.847%. We saw that the actual proportion estimates of donors hovered around the expected average proportion. Furthermore, each individual’s estimated proportion was tightly distributed within same-day replicates - the standard deviation of individual’s proportion estimates across the same-day replicates was never greater than 0.237% (**Figure 4**). We found the proportion estimates were very consistent across same-day replicates, with a Pearson’s correlation coefficient (*r*) greater than 78% for each individual cell line’s estimated proportion across every within-experiment comparison.

**Figure 4:**
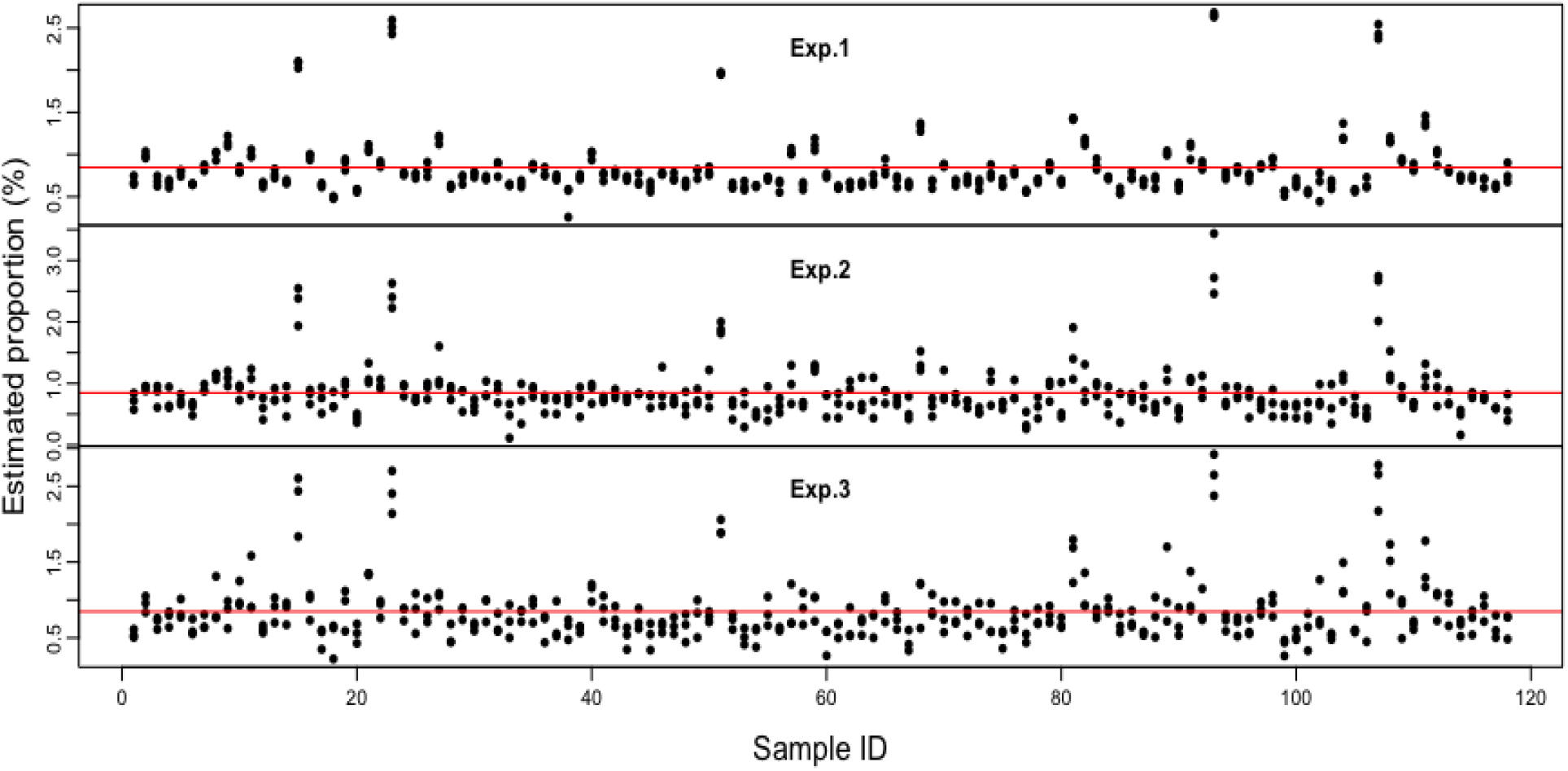
Spread of same-day replicate estimated proportions within Exp.1, Exp.2, and Exp.3 The figure shows the spread of estimated proportions for each individual within the same-day replicates of Exp.1, Exp.2, and Exp.3. Sample IDs (x-axis) are plotted against the estimated proportions (y-axis). Each unique sample has 3 estimated proportions plotted on the y-axis, representing that individual’s estimated proportion from each replicate. We see that individual cell lines’ proportions are tightly distributed within replicates, and the estimated proportions hover around the red horizontal line, which corresponds to the expected proportion for each individual, 0.847%.

When comparing proportions across different day experiments, the results also show a very strong positive correlation of an individual’s average proportion in experiments 1, 2, and 3 (*r*>0.94, *P*<2.2 × 10^−16^ across each comparison) (**Figure 5**). This result suggests that an individual’s reprogramming ability is consistent both within a single day and across multiple experimental runs separated by several weeks.

**Figure 5:**
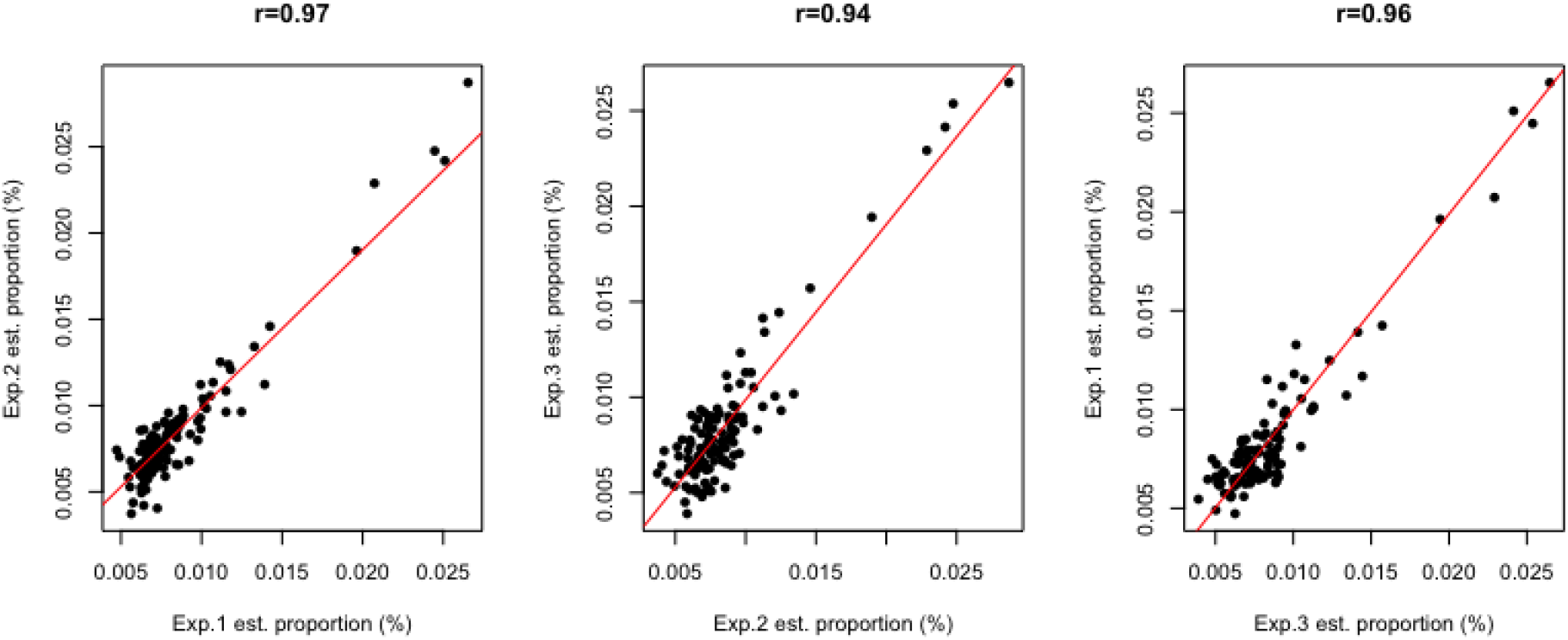
Average donor proportion estimate correlations across Exp.1, Exp.2, and Exp.3 The figure shows the cross-experiment correlations of individual cell lines’ proportions in the pooled cohort. We see that individual cell lines’ proportions stay quite consistent across experiments, with a Pearson correlation coefficient greater than or equal to 0.94 across each comparison.

### Genetic basis of successful reprogramming ability

#### Assessment of SNPs and heritability estimates associated with reprogramming ability

Before constructing our reprogramming ability test statistic, we excluded two out of the 118 individuals in our final cohort that were stratified during PCA (See Materials and Methods).To estimate reprogramming ability for each donor, we calculated the average estimated proportion for each individual across our three experimental runs and then performed a rank-based inverse normal transformation of the averages to obtain a reprogramming ability Z-score for each individual in the purified iPSC pool. We did not identify any genome-wide significant SNPs (*P*<5×10^−8^) associated with reprogramming ability (**Figure 6**).

**Figure 6:**
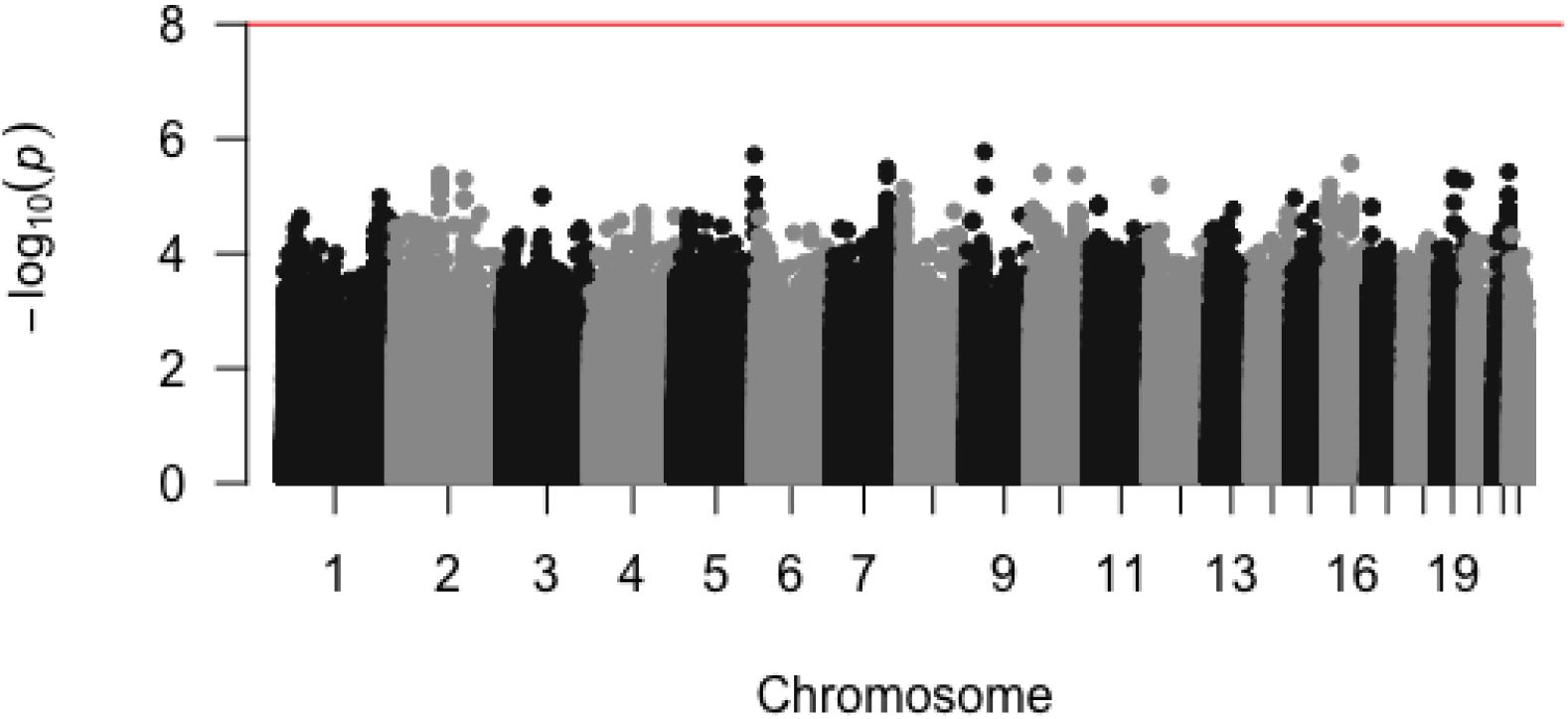
Manhattan plot for our genome-wide association study of reprogramming ability This figure shows the association of SNP genotype and reprogramming ability was evaluated using a univariate linear regression model for 2,485,057 SNPs in the discovery GWAS of the 118 successfully reprogrammed donor lines in our pooled cohort. P-values (-log_10_(*P*), y-axis) were plotted against the respective chromosomal position of each SNP (x-axis). No SNP achieved genome-wide significance (*P*<5×10^−8^), marked in red.

We used GCTA LD-score regression to estimate the proportion of variance explained by common SNPs (heritability, or h^2^_g_) for reprogramming ability(Yang et al., 2011). Given the low sample size, we did not obtain a meaningful heritability estimate for reprogramming ability (h^2^=1.0±0.47). The combination of lack of genome-wide significant SNPs and inconclusive heritability estimates suggests low power due to small sample size and limited ability to explain heritability of both reprogramming ability from associated loci.

#### Assessment of reprogramming ability with age and sex of donor cells

Of the 118 successfully reprogrammed individuals in our pooled cohort, there were 82 males and 36 females, ranging in age from 23 to 90 years old at the time of sample collection (**Table S3**). We used simple linear regression to look for association between both age and sex with reprogramming ability. There was no significant association between donor age and reprogramming ability Z-score (R^2^=3.93×10^−4^, *P*=0.833) (**Figure 7A**), nor any significant association between sex and reprogramming ability Z-score (R^2^=0.0140, *P*=0.105). (**Figure 7B**).

**Figure 7:**
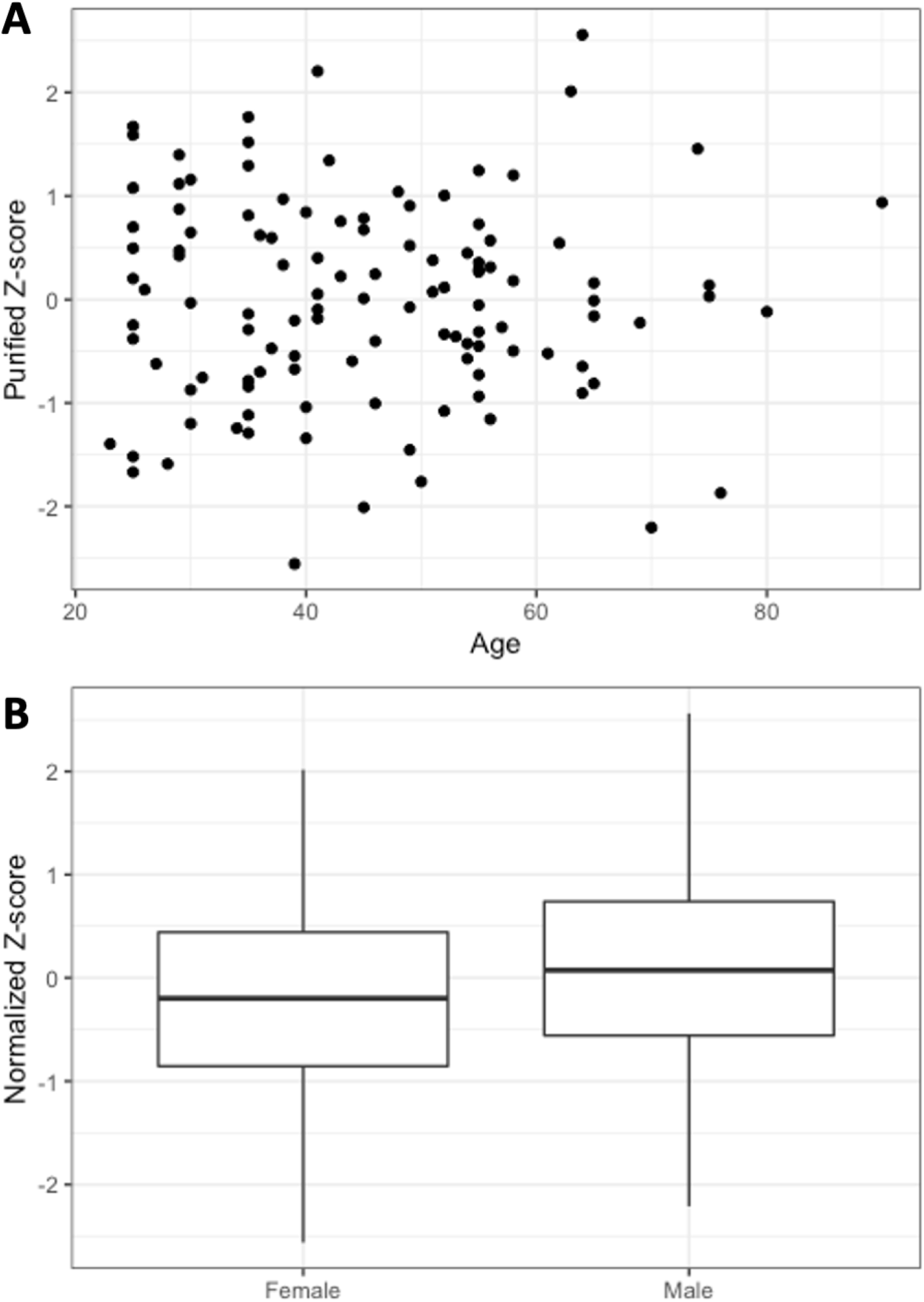
Plots showing sample age and sex versus reprogramming ability Z-score (**A**) Each individual’s age plotted against their reprogramming ability Z-score. Each individual sample’s age (x-axis) was plotted against their reprogramming ability Z-score (y-axis). No significant association was found between age and reprogramming Z-score (R^2^=3.93×10^−4^, *P*=0.833). (**B**) Box-and-whisker plots comparing male versus female reprogramming ability Z-scores in the unpurified iPSC populations. The x-axis reflects the male versus female population and the y-axis reflects normalized Z-scores. There was no significant association between sex and reprogramming (R^2^=0.0140, *P*=0.105).

### Expression of immunoglobulin lambda variable precursor genes significantly associated with both reprogramming success and ability

Using gene expression data obtained from RNA sequencing data for LCLs (See Materials and Methods), we checked for association between gene expression and reprogramming ability across 58 different LCL lines present in the pool of 120 donors. Of the 58 lines, one individual was stratified from the central population cluster after performing PCA. Removing this individual from gene expression analysis did not result in measurable change in either the p-values or false discovery rates of genes with expression significantly associated with reprogramming ability.

After grouping individuals by their reprogramming ability Z-score and fitting genewise negative binomial generalized linear models in edgeR(Robinson et al., 2010), we examined the association of gene expression with reprogramming ability (**Table S5)**. Five genes showed expression significantly associated with change reprogramming ability: IGLV10-34 (*P*=3.33×10^−7^, FDR=1.73×10^−3^), IGLV3-9 (*P*=8.30×10^−7^, FDR=2.16×10^−3^), IGLV1-17 (*P*=1.36×10^−6^, FDR=2.36×10^−1^), IGLV1-6 (*P*=5.19×10^−6^, FDR=6.74×10^−3^), and IGLV3-1 (*P*=2.43×10^−5^, FDR=0.025) These are protein-coding genes for the variable domain of immunoglobulin light chains that participate in antigen recognition(Collins and Watson, 2018). This indicates that pre-reprogramming expression of immunoglobulin precursor genes might be associated with increased reprogramming ability in LCLs.

**Table 1:** LCL gene expression association with reprogramming ability

|  | logFC | logCPM | F | PValue | FDR |
| --- | --- | --- | --- | --- | --- |
| IGHV4-34 | -1.3785791 | 6.284607 | 33.1836 | 3.33E-07 | 0.001734632 |
| IGKV1-9 | 1.3891019 | 5.385836 | 30.44276 | 8.30E-07 | 0.002158794 |
| IGKV1-17 | -1.6585554 | 7.873226 | 28.98731 | 1.36E-06 | 0.002364993 |
| IGKV1-6 | 1.3468997 | 5.064504 | 25.20039 | 5.19E-06 | 0.00674652 |
| IGLV3-1 | 1.864739 | 7.550449 | 21.05404 | 2.43E-05 | 0.025231739 |
| IGLV1-51 | 1.3129681 | 5.650807 | 14.82982 | 2.95E-04 | 0.255971371 |
| IGLV6-57 | -0.9073692 | 4.582162 | 14.2343 | 3.80E-04 | 0.282448173 |
| IGHA2 | 0.7529145 | 5.467972 | 10.95823 | 1.60E-03 | 0.918921186 |
| TRNC | 0.2082136 | 7.549076 | 10.79156 | 1.73E-03 | 0.918921186 |
| TRNA | 0.2808821 | 4.955505 | 10.7448 | 1.77E-03 | 0.918921186 |

This table shows the results of the top ten genes with expression from LCLs most associated with reprogramming Z-scores grouped by quintiles. Gene expression data was obtained through edgeR analysis using RNA-seq expression profiles of 58 individuals from our cohort of 118 successfully reprogrammed pooled lines.

## Discussion

In this study we successfully demonstrate the ability to generate a valuable research resource for scientific studies. Our method for induction of pluripotency from pooled cells, iPPC, produces a line of pooled iPSC which contains cells from over 100 genetically distinct donor lines. Using a deconvolution algorithm based on whole-genome sequencing (WGS), the proportional presence of each donor is attained and can be used as a basis for downstream analysis and assays. We observed a yield of 98%, with 118 of 120 donors being present in our final iPSC lines after a 5.5 week experimental protocol and proportion estimates tightly distributed around the expected average in three separate experiments. We additionally found strong correlations between our same-day replicates and experiments. Overall, iPPC demonstrates a reliable and time efficient method for studying large cohorts of donors from a variety of genetic backgrounds.

The high correlation found between same-day replicates, *r*>0.78, shows that the estimated donor proportions are not the result of stochastic reprogramming events and the similarly high correlation between different-day experimental replicates, *r*>0.94, indicates variability introduced by pooling, culturing or nucleofection does not affect eventual proportions. This suggests an inherent element to a cell’s ability to reprogram into iPSCs and that our final proportions can be used as a proxy for donor reprogramming ability. Looking for population influences, we conducted association analyses for donor age and sex. While we did not discover significant correlations or associations from our limited sample size, iPPC enables the expansion to much larger sample sizes for similar genetic analysis.

iPSC phenotypes such as RNA expression, differentiation potential, cell behavior, and chromosome stability have been linked to genetic differences between donor lines(Thomas et al., 2015; Kyttälä et al., 2016; Kilpinen et al., 2017; DeBoever et al., 2017; Panopoulos et al., 2017; Banovich et al., 2018; Vigilante et al., 2019; Mirauta et al., 2020; Cuomo et al., 2020). While our study found no SNPs significantly associated with reprogramming ability, the cell lines we have generated through iPPC are a useful tool when further investigating the impact of genetics on these iPSC phenotypes. We can evaluate differences with a significant reduction in background signal that typically results from individual reprogramming and culture. Additionally, the lack of association in our data may have resulted from our small sample size, with 118 donors not being sufficient for detecting genetic influences on reprogramming ability. Other potential issues of iPPC include the effect of genetic aberrations as a result of the reprogramming process(Araki et al., 2020; D’Antonio et al., 2018). This can be resolved by performing downstream clonal selection, donor identification and subsequent quality control to filter out aberrant iPSC clones(Chen et al., 2021).

Having established that the final iPPC proportions are non-random, but finding no genetic basis, we looked at the influence of pre-reprogramming cell states and found five significantly associated genes all belonging to the V region of the variable domain of immunoglobulin light chains. These associated genes are possibly linked to our cell type of origin, LCLs. Originating from Epstein Barr Virus (EBV) infection of B cells from peripheral blood(Mrozek-Gorska et al., 2019), this association possibly results from an overall increase in expression of immunoglobulin precursor genes. This suggests further investigation into whether genetic perturbation of these genes can be the basis of improving iPSC reprogramming efficiency of LCLs.

The use of iPPC will result in a paradigm shift towards performing *in-vitro* genetic studies. Instead of testing the phenotypes of each donor cell independently, iPPC suggests a pooled approach as a way to multiplex the experiment. iPPC generates a pool of multi-donor iPSCs, which can lead to several potential downstream research applications (**Figure 8**). The pooled multi-donor iPSCs can be differentiated into a wide variety of adult tissue types for genetic testing to determine if there are any genetic variant associations with *in-vitro* phenotypes.

**Figure 8:**
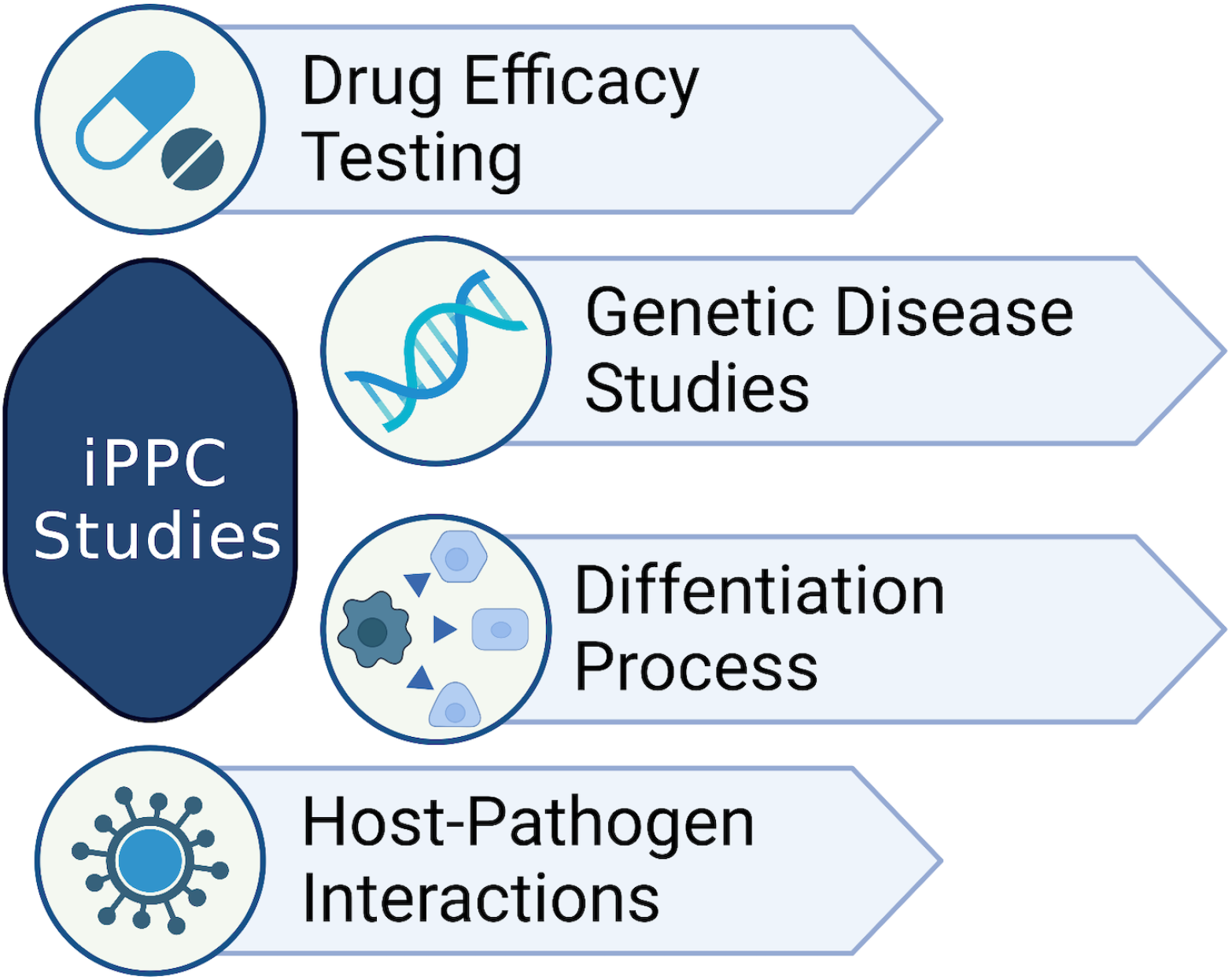
Potential downstream research applications of IPPC Some examples of the use of IPPC for research.1) IPPC can be used for studying drug efficacy by identifying donors there are more or less susceptible to the effect of the drug. 2) IPPC can be used for studying human genetic diseases by testing the association of genetic variants with *in-vitro* phenotypes. 3) IPPC can be used for studying the effect of different genetic backgrounds on specific biological processes, e.g. differentiation. 4) IPPC can be used for studying host-pathogen interactions by identifying donors that are more or less susceptible to pathogenic infections.

Our experimental method shows promise for its utility in an assortment of experiments. Providing researchers with a robust method for interrogating underlying factors influencing iPSC technology with a minimization of experimental background, iPPC is a practical and scalable approach to helping researchers expand their studies and sample sizes. Utilizing other cohorts, such as the 1000 Genomes Project(1000 Genomes Project Consortium et al., 2015) which contains over 2000 donor lines of multiple ancestries with WGS data, we can significantly expand the power of future studies. Pooled iPSC lines can be used in conjunction with a variety of other experimental techniques such as single cell sequencing, GWAS, differentiation assays amongst others. With WGS of donor lines, researchers can conduct studies of patient derived iPSCs effectively. We are confident our iPPC method can be expanded to a variety of applications which will enrich the quality of future scientific inquiry.

## Materials and Methods

### Cohort Description

We obtained 120 Harvard Personal Genome Project (PGP) unique lymphoblastoid donor cell lines (LCLs) from the Coriell Institute for Medical Research (https://coriell.org/)(Ball et al., 2012, 2014; Chan et al., 2017; Mao et al., 2016). These PGP cell lines come with whole-genome sequencing genotype data as well as selected phenotypes associated with each donor (**Table S1**)(Ball et al., 2012, 2014; Chan et al., 2017; Mao et al., 2016).

### Lymphoblastoid cell lines

#### LCL Culture

Lymphoblastoid cell lines (LCLs) were cultured in RPMI media with 10% Fetal Bovine Serum (FBS) and 1% Pen Strep, Gibco 72400-047, 10438-026, 15140-122, in a 37°C, 5.0% CO_2_ incubator. Cultures were maintained in 3ml of media in 12-well culture plates with one cell line per well. Media was changed every 3-4 days by replacing 2/3 of old media and cells were passaged as needed by resuspending the culture and discarding 2/3 of suspension before replacing the media. Before every media change, the health of each cell line was visually evaluated.

#### LCL Thawing

LCLs were kept frozen under -80°C or LN2 conditions in vials containing 3-5×10^6^ cells in 1ml of 95% FBS and 5% DMSO. Vials were thawed by placing them in a 37°C water bath until only a small piece of ice remained. The vials were put on ice until being gently transferred to 15ml conical tubes. 9ml of cold RPMI media was slowly dripped onto the cells before being gently mixed by inversion. Cells were then spun down at 300 x g for 5min at 4°C. The supernatant was poured off, and the cells were resuspended in 12ml of culture media before being transferred to a 25cm^3^ flask and grown in a 37°C, 5.0% CO_2_ incubator. After 1 week, 1×10^6^ cells were transferred to a single well on a 12-well plate for experiments while the remaining cells were cultured further before being frozen.

### Induction of Pluripotency form Pooled Cells, iPPC

#### Cell Counting

On day 0 (D=0), LCLs were resuspended by pipetting. 200ul of suspended culture was transferred to a 96 v-bottom plate. LCLs were then counted using a MACSquant VYB flow cytometry machine. Using a 25ul uptake volume, the cell count was acquired through gating the live cell population and recording the cell count calculated by the software. Cell counts were recorded manually. The count was used for pooling the different cell lines in equal proportion.

#### Nucleofection/Culturing

At day 0 (D=0), cell lines were pooled evenly such that the total cell count was 4 × 10^7^ cells. The pool of cells was split into 4 replicates of 1 × 10^7^ cells. The other three replicates were nucleofected with reprogramming plasmids gCXLE-hOCT3_4shp53-F, gCXLE-hSK, and gCXLE-hUL (Addgene #27077, #27078, #27080 respectively). These plasmids expressed hSOX2, hKLF4, OCT3/4, c-myc, hLIN28, and shp53. The SF cell line solution was prepared fresh by combining 18.2ul of SF supplement, 81.2ul SF solution and 7.5ug plasmid per nucleofection, Lonza V4XC-2024. The reprogramming used 2.5ug of OCT3/4, HSK, and HUL each per replicate. The pooled 1 × 10^7^ cells were spun down in 1.5ml tubes. The supernatant was completely removed from the pelleted cells. The pellets were resuspended in 100ul of the prepared nucleofection solution and immediately transferred to the nucleovettes. Nucelovettes were placed in the Lonza 4D Nucleofector and run using the K562 cell-line setting with SF solution. After nucleofection, cells were gently transferred to prepared wells containing 1ml of warm RPMI and incubated for 18 hours at 37°C.

At D=1, RPMI was removed from the pooled LCL replicates nucleofected with reprogramming plasmid and replaced with ReproTeSR Medium (Stem Cell #05919). Cells were then transferred to 6-well culture plates coated with 8.6ug/cm^2^ Matrigel. Each replicate being placed in 2 wells. Half the ReproTeSR volume was exchanged for fresh media every other day until D=13.

### iPSC Culturing

On D=13 unsuccessfully reprogrammed LCLs remained in suspension and were removed from the wells by washing twice with DPBS before replacing the media with StemFlex Basal Medium (ThermoScientific A33493-01). The remaining adherent cells were detached from the wells using Accutase and plated on Matrigel coated 75cm^2^ flasks with StemFlex and 10μM of Y-27632 dihydrochloride (Abcam ab120129). After 24 hours, we replaced the media for StemFlex without Y-27632 dihydrochloride.

Once the flasks reached 90% confluency, samples of cells were collected for DNA extraction and flow cytometry analysis prior to magnetic column purification. 1×10^7^ cells per replicate were purified using 100ul of Pluripotent Stem Cell microbeads (Miltenyi Biotec 130-095-804), and LS columns (Miltenyi Biotec 130-042-401). The isolated iPSCs were again plated on Matrigel Matrix, REF#354234, coated 75cm^2^ culture flasks. When the purified iPSC replicates achieved 90% confluency, samples were again collected for DNA extraction and flow cytometry analysis. Remaining cells were frozen and stored at -80°C in aliquots of 1×10^7^ cells in 1ml of 95% FBS and 5% DMSO freezing media.

### Flow Cytometry Analysis

iPSC samples were prepared by detaching them from Matrigel coated wells using Accutase. Detached cells were counted using a NanoEnTek EVE automated cell counter, and 200,000 cells were transferred to wells in a 96 V-bottom plate. Cells were then pelleted and resuspended in 50ul of DPBS with 1:50 of each antibody for 1 hour at 4°C along with unstained iPSC and HEK293T controls. Antibodies used were TRA-1-60 and SSEA-4 both conjugated with Alexa Fluor 488 (Novus Biologicals NB100-730AF488 and NBP2-26644AF488 respectively). After staining, cells were washed 3 times with DPBS, then resuspended in 100ul DPBS for flow analysis.

Samples were run on a MCASQuant VYB instrument with an uptake volume of 50ul. Positively stained cells were determined by first gating the live cell population on FCS/SSC channel and then gating 488 positive signals on B1/SSC using unstained control.

### Next-generation Sequencing of DNA

Genomic DNA was extracted using BIONEER AccuPrep Genomic DNA Extraction Kit, Ref#K-3032 from a sample of 1×10^6^ cells per replicate. A DNA library was prepared from these samples using the Invitrogen Colibri PCR-free ES DNA Library Kit, Ref# A38577096.

The prepared DNA libraries were then quantified using the Fragment Analyzer Service of the UMass MBCL core facility (https://www.umassmed.edu/nemo/mbcl/fragment-analyzer-service/) which uses the Advanced Analytical (Agilent) capillary electrophoresis system. The DNA libraries were then pooled and sent for whole-genome sequencing (Psomagen). We performed Illumina paired-end sequencing (read 150 bases) on the HiSeq X Ten system, with different index barcodes for each sample resulting in roughly one million paired-end reads per sample on average (**Table S4**).

### RNA sequencing of LCLs and whole transcriptomic profiling

For a subset of donor LCLs from the PGP, we have RNA sequencing data available that was generated from a different project (https://www.synapse.org/#!Synapse:syn25927432). The RNA sequencing was obtained by first culturing the LCLs overnight (2×10^6^ cells per donor in 1 well of a 6 well plate) in RPMI medium supplemented with 10% Fetal Bovine Serum (FBS) and 1% penicillin-streptomycin (2ml of media per well of a 6-well plate). The following day, the LCLs were harvested and we performed total RNA extraction of the LCLs (Invitrogen PureLink RNA Mini Kit). We then sent the extracted RNA for whole transcriptome sequencing (Psomagen).

Preparation of samples for RNA-Seq analysis was performed using the TruSeq RNA Sample Preparation Kit (Illumina, San Diego, CA). Briefly, rRNA was depleted from total RNA using the Ribo-Zero rRNA Removal Kit (Human/Mouse/Rat) (Illumina, San Diego, CA) to enrich for coding RNA and long non-coding RNA, following the TruSeq Stranded Total RNA Sample Prep Guide, Part # 15031048 Rev. The Ribo-Zero libraries were sequenced on the Illumina NovaSeq 6000 System with 151 nucleotide paired end reads, according to the standard manufacturer’s protocol (Illumina, San Diego, CA).

The raw sequence reads were aligned to human genome hg38 (ensembl_GRCh38, GenBank Assembly ID GCA_000001405.15) with the star aligner (v2.5.2b)(Dobin et al., 2013) and gene level expression (read counts) were summarized by the “--quantMode GeneCounts” parameter in star.

### Estimating donor proportion

We performed alignment of the sequencing reads for each sample to the GRCh37 (hg19) assembly of the human genome using bwa (version: 0.7.17-r1188)(Li and Durbin, 2009). We obtained whole-genome SNP genotyping data for each donor from the Harvard Personal Genome Project (PGP) website (https://pgp.med.harvard.edu/). We interrogated a list of 9,923,808 autosomal SNP positions identified within the PGP data for the aligned reads using samtools(Li et al., 2009) mpileup (version 1.9) and determined that 500 thousand to 8.1 million SNPs are aligned to at least 1 sequencing read within each of the pooled samples (**Table S5**)(Li et al., 2009). We then formatted the known alleles for data obtained from each of the pooled samples and performed the donor proportion estimation using the PoolSeq program after running for 2000 iterations (https://pgpresearch.med.harvard.edu/poolseq)(Chan et al., 2018). Estimates from the final iteration are then used as the donor proportion for each sample.

### Determining presence or absence of donor cells within pools

For the original cohort, we modeled the donor proportion of absent donors (null distribution) using the proportions obtained from the 20 individuals not present in the pools (**Table S1**). The mean ± standard deviation estimates of the proportions were 0.0003884938557 ± 0.0002337132416 in Exp. 1, 0.0004763303045 ± 0.0005720989657 in Exp. 2, and 0.0003598711056 ± 0.0005713536307 in Exp. 3. We used a stringent cutoff of 0.0028 such that proportions above this cutoff would be considered present while proportions below this cutoff would be considered absent. Using this cutoff would give a false discovery rate (FDR) of 1.43 × 10^−6^. This cutoff resulted in a pool of 118 donors remaining from the initial 120 donors.

### Quality control and association analysis

Plink 1.90b6.9(Chang et al., 2015) was used to perform quality control and association analysis. We ensured that no individuals had a genotyping rate less than 20%, discordant sex data, or were duplicated. SNPs with minor allele frequency less than or equal to 0.02, as well as those not in Hardy-Weinburg equilibrium (*P*<1.0×10^−6^) were removed. Of the 118 samples in the final pool of donors, we excluded two individuals of non-European ancestry that were stratified in PCA. (**Figure S1**). We used Plink to implement a univariate linear regression model for 2,485,057 SNPs in the discovery GWAS of the 118 successfully reprogrammed donor lines in our pooled cohort.

## Acknowledgements

We thank the Harvard Personal Genome Project (PGP) and the Coriell Institute for Medical Research (CIMR) for contributing and making the cell lines available for research.

## Ethics Declaration

### Ethics approval

The use of the human cell lines for research in this work was determined to not be human subjects research by the IRB of UMass Chan Medical School (H00021419).

---

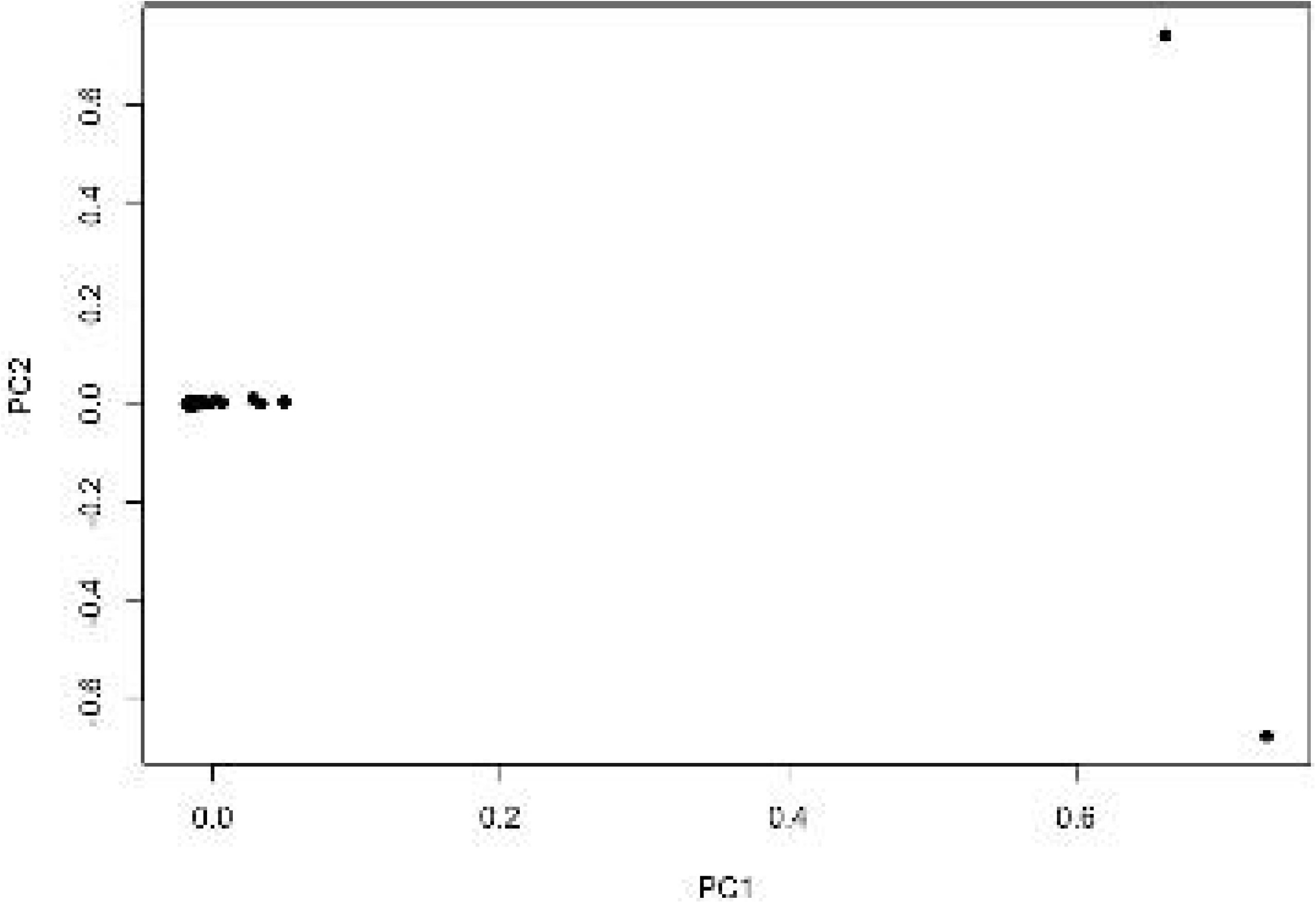

